# Simultaneous 3D Cellular Positioning and Apical Dendritic Morphology of Transgenic Fluorescent Mouse CA3 Hippocampal Pyramidal Neurons

**DOI:** 10.1101/2022.08.12.503761

**Authors:** Christopher J. Handwerk, Katherine M. Bland, Collin J. Denzler, Anna R. Kalinowski, Cooper A. Brett, Brian D. Swinehart, Hilda V. Rodriguez, Hollyn N. Cook, Elizabeth C. Vinson, Madison E. Florenz, George S. Vidal

**Affiliations:** Department of Biology, James Madison University, MSC 7801, Harrisonburg, Virginia 22807, United States of America

**Keywords:** CA3, pyramidal neuron, dendritic morphology, dendritic complexity, reconstruction, GFP-M

## Abstract

**Background:** Pyramidal neurons throughout hippocampal CA3 are diverse in their dendritic morphology, and CA3 is not homogenous in its structure or function. Nonetheless, few structural studies have captured the precise 3D somatic position and the 3D dendritic morphology of CA3 pyramidal neurons simultaneously.

**New method:** Here, we present a simple approach to reconstruct the apical dendritic morphology of CA3 pyramidal neurons using the transgenic fluorescent *Thy1*-GFP-M line. The approach simultaneously tracks the dorsoventral, tangential, and radial positions of reconstructed neurons within the hippocampus. It is especially designed for use with transgenic fluorescent mouse lines, which are commonly used in genetic studies of neuronal morphology and development.

**Results:** We demonstrate how topographic and morphological data are captured from transgenic fluorescent mouse CA3 pyramidal neurons.

**Comparison with existing methods:** There is no need to select and label CA3 pyramidal neurons with the transgenic fluorescent *Thy1*-GFP-M line. By taking transverse (not coronal) serial sections, we preserve fine dorsoventral, tangential, and radial somatic positioning of 3D-reconstructed neurons. Because CA2 is well defined by PCP4 immunohistochemistry, we use that technique here to to increase precision in defining tangential position along CA3.

**Conclusions:** We developed a method for simultaneously collecting precise somatic positioning as well as 3D morphological data among transgenic fluorescent mouse hippocampal pyramidal neurons. This fluorescent method should be compatible with many other transgenic fluorescent reporter lines and immunohistochemical methods, facilitating the capture of topographic and morphological data from a wide variety of genetic experiments in mouse hippocampus.

**Highlights:**

- Simultaneous capture of 3D location and pyramidal dendritic morphology in CA3
- Method utilizes replicable techniques and reagents available to most laboratories
- Method is adaptable to other transgenic mouse lines and immunohistochemical studies

## 1 Introduction

Hippocampal pyramidal neurons are variable in their gene expression, structure, and function^1–4^. Some heterogeneous structural and functional features of these neurons vary in a spatially patterned way. For example, in CA3, place field size, pattern completion, and sharp wave initiation are all arranged topographically^3,5–9^. Specific locations within CA3 are recruited during both spatial^10–14^ and non-spatial learning and memory^15^. Focal lesions within specific CA3 subfields can lead to disrupted fear memory^16^ or spatial processing^17^. Thus, for studies of CA3 pyramidal dendritic morphology, it is critical to control for the precise topographic location of analyzed neurons.

Prior morphological studies can estimate the topographic location of CA3 somata via *post hoc* analysis. In other words, the somatic position of a neuron in an experimental section can be estimated by approximating the section’s topographic characteristics to a map or atlas. We developed a method to capture both the precise 3D location of a neuron and its 3D dendritic morphology simultaneously. We do so by focusing on the three axes of the sheet-like *stratum pyramidale* of CA3, home to CA3 pyramidal neuron somata. One axis is tangential, where neurons are more proximal or more distal to the dentate gyrus. Another axis is dorsoventral (or “long”), where neurons are closer or further away from the septum. A third axis is radial, where neurons are deep or superficial within the *stratum pyramidale*.

In designing our new method, we sought to rely on relatively simple laboratory techniques to increase its reproducibility and access by other laboratories. Our prior work used serial sections and sparsely labeled, green fluorescent protein (GFP)-filled neurons to discover a gradient of dendritic morphology across layer 2/3 of the mouse cerebral cortex^18^, which is perturbed by the deletion of integrin β3 (*Itgb3*), an identified autism risk gene^19^. Here, we applied the same simple serial sectioning strategy, this time pairing it with the widely available and characterized *Thy1*-GFP-M line^20^ as a relatively easy way to fill sparse subsets of CA3 pyramidal neurons with GFP.

This fluorescent method should be compatible with many other transgenic fluorescent reporter lines and immunohistochemical techniques, facilitating the capture of topographic data (*i.e*., 3D somatic position) and morphological data (*i.e*., 3D apical dendritic reconstruction) from a wide variety of genetic experiments in mouse hippocampus.

## 2 Material and methods

A graphical flowchart of the main steps of the method is presented in Figure 1, and the details for each step are presented below.

**Figure 1.**
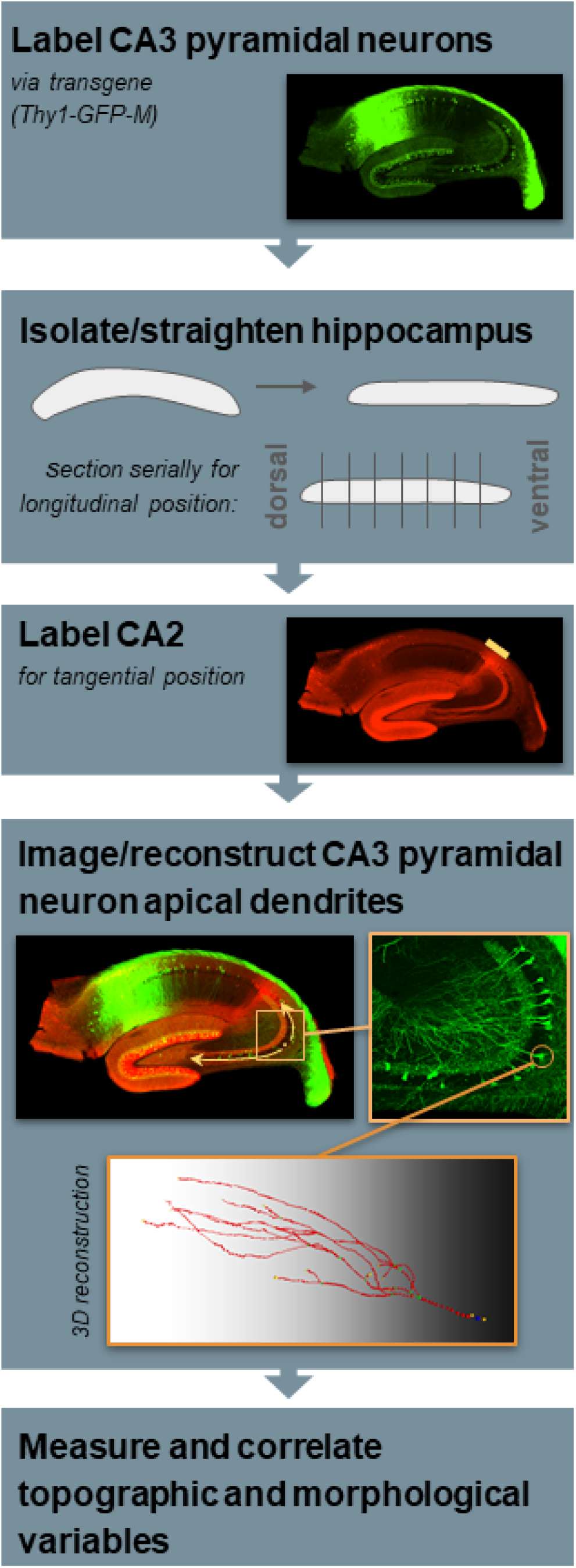
Schematic flowchart of methods.

### 2.1 Ethics statement

Our study complies with the ARRIVE 2.0 guidelines by including essential information regarding study design, sample size, inclusion/exclusion criteria, randomization, blinding, outcome measures, statistical methods, experimental animals, experimental procedures, and results, where applicable. The James Madison University Institutional Animal Care and Use Committee approved the necessary work with mice (protocol #20-1067). We followed the guidelines of the National Research Council’s Guide for the Care and Use of Laboratory Animals.

### 2.2 Mice

In our mouse colony, the *Thy1*-GFP-M transgenic mouse^20^ (STOCK Tg(Thy1-EGFP)MJrs/J, The Jackson Laboratory #007788) had been backcrossed to the C57BL6/J strain (The Jackson Laboratory #000664) for at least 5 generations by the time we bred mice for this study. Male transgenic (*i.e*., hemizygous) *Thy1*-GFP-M mice were bred with female mice without the transgene to produce experimental mice. Approximately half of the progeny from this cross carried the transgene, as determined by genotyping of tissue acquired via ear punch (Transnetyx; probe “Thy1-3 Tg”), and as reported previously.^21^ We selected two hemizygous mice at postnatal day 30 for the experiment. Selected mice were female by chance, and we did not attempt to measure the estrus cycle of the mice at this age. This study is therefore unable to assess any potential differences between male and female mice, or any potential effect of estrus cycle on our results. Selected mice were not visibly abnormal in weight.

After birth, mice in our colony are housed with their parents and littermates until weaning. Mice are weaned approximately between postnatal day 21 and 28. Upon weaning, mice are housed with their same-sex littermates. All mice are housed in a temperature- and humidity-controlled, specific-pathogen-free environment with a 12-h light/dark cycle. Mouse cages are supplied with Teklad ¼” corncob bedding, with food (Teklad 18% protein rodent diet) and water supplied *ad libitum*.

### 2.3 Histology

#### 2.3.1 Perfusion and dissection

We euthanized the mice by intraperitoneal injection of ketamine (240 mg/kg)-xylazine (48 mg/kg)-acepromazine (1.85 mg/kg). Immediately after cutting through the thoracic wall, we transcardially perfused approximately 50 mL of ice-cold 1× phosphate-buffered saline (PBS; diluted from OmniPur® 10× PBS Liquid Concentrate, Sigma #6505, with Milli-Q distilled water) for 5-10 minutes, immediately followed by approximately 50 mL of ice-cold, freshly made 4% (wt/vol) paraformaldehyde (Sigma-Aldrich) in PBS, pH 7.4 (4% PFA) for 5-10 minutes. Note that this method of perfusion may have some inherent disadvantages due to the 5-10 minutes in between the opening of the thoracic wall and the fixation of the brain (see Discussion).^22^ We dissected the head of each mouse and post-fixed it in 4% PFA for approximately 24 h at 4 °C. We dissected the brain out of the skull using two #5SF forceps (Fine Science Tools #11252-00) and a micro spatula (Fine Science Tools #10091-12) and dissected away the meninges. We made a midsagittal cut with a razor blade to separate the hemispheres. Keeping the brain moist with 1× PBS, we scooped out subcortical structures from each hemisphere (for this study, the left hemisphere) with the micro spatula, leaving the cortex and hippocampus intact. After removing as much white matter as possible near the hippocampus proper, we used the micro spatula to loosen the hippocampus proper from the cortex, then used the forceps to severe the hippocampus proper from the cortex. We isolated 4 hippocampi from the 2 mice (2 left and 2 right).

#### 2.3.2 Straightening the hippocampus

We placed each hippocampus on the middle of a glass slide, with the dorsal and ventral ends pointing toward the top and bottom of the slide. At this point in the procedure, the hippocampus still exhibits a curvature along the dorsoventral axis. We adapted prior procedures that provide and discuss methods to take exact transverse sections by straightening the dorsoventral axis^23,24^. We stacked 22 × 22 mm #1 coverslips to match the height of the hippocampus (approximately eight coverslips) and placed this stack to the left of the hippocampus. We stacked the same number of 22 × 22 mm #1 coverslips to the right of the hippocampus. Then, we gently pushed the two coverslip stacks together, straightening the hippocampus. We placed a second glass slide on top and held the entire configuration together by binding the glass slides together with mini binder clips. We immersed the entire configuration in a Petri dish with 1% PFA (wt/vol) diluted in 1× PBS and incubated it overnight at 4 °C.

#### 2.3.3 Sectioning the hippocampus

We embedded each hippocampus in agar prior to sectioning. To do so, we first dissolved and melted 4% agar (wt/vol, in Milli-Q distilled water) on a hot plate. Off the heat, we stirred the agar as it cooled. Just before it hardened, we poured a 3-5 mm base of molten agar into a mold (2.5 mL glass micro beaker, Electron Microscopy Services #60982), laid the straightened hippocampus on top, and poured more molten agar on top of the hippocampus. Once the agar hardened, we blocked the agar on the platform of a vibrating microtome (Leica VT1000S), such that we could take 100 μm-thick free-floating transverse sections in 1× PBS. Critically, we collected each section in order on a 96-well, round-bottom micro plate that was pre-loaded with 1× PBS. We found that removing the agar around each free-floating section was not strictly necessary, but greatly aided in the pipetting that is required for subsequent immunohistochemistry (see below).

#### 2.3.4 Immunohistochemistry

To enhance GFP fluorescence, we used the Mouse on Mouse (M.O.M.) Immunodetection Kit (Vector Laboratories, #BMK2202) in conjunction with mouse monoclonal anti-GFP (clone “3E6”, Invitrogen #A-11120), and goat anti-mouse IgG (H+L) F(ab’)_2_ fragment conjugated to Alexa Fluor 488 (Cell Signaling Technology, #4408). Each well, containing one section, received 50 μL of solution at each step. We followed the manufacturer’s instructions for preparing Mouse on Mouse (M.O.M.) Immunodetection Kit blocking solution and diluent. The ingredients of both the blocking solution and diluent are provided with the kit and are proprietary.

We incubated the tissue in blocking solution for 1 h, washed twice in 1× PBS, incubated in diluent for 5 minutes, incubated in primary antibody (1:1000 in diluent) for 30’, rinsed twice in 1× PBS, incubated in secondary antibody (1:250 in diluent), and rinsed thrice in in 1× PBS.

To stain for CA2 pyramidal neurons, we used rabbit polyclonal anti-PCP4 (Invitrogen #PA5-52209) and highly cross-adsorbed goat anti-rabbit IgG (H+L) conjugated to Alexa Fluor 594 (Invitrogen #A-11037). Each well, containing one section, received 50 μL of solution at each step. We incubated the tissue in blocking solution (5% [vol/vol] normal goat serum [Vector Laboratories #S-1000-20] and 0.15% [vol/vol] Triton X-100 [Sigma-Aldrich #T8787] in 1× PBS), incubated in primary antibody (1:1000 in blocking solution), rinsed twice in 1× PBS, incubated in secondary antibody (1:600 in blocking solution), and rinsed twice in 1× PBS.

#### 2.3.5 Mounting and storing slides

We mounted sections directly on charged glass slides (Globe Scientific #1358L), applied ProLong Diamond Antifade Mountant (Invitrogen #P36970), and placed a #1 coverslip (Globe Scientific #1315-10) on top. We cured the slides for at least 48 h before imaging. For long-term storage, we have kept the slides at −20 °C.

### 2.4 Microscopy

#### 2.4.1 Hippocampal sections

We took images of each hippocampal section to identify somata with GFP and/or PCP4 expression, using a Nikon Eclipse Ti-2 microscope with a 4× PlanApo objective (0.20 NA) in the “green” channel (excitation filter: 446-486 nm, emission filter: 500-550 nm, to capture GFP and Alexa Fluor 488 signal), and in “red” (excitation filter: 542-582 nm, emission filter: 604-678 nm, to capture Alexa Fluor 594 signal). We imaged 80 sections (18, 16, 24, and 22 sections from hippocampi 1-4, respectively) for this study.

#### 2.4.2 CA3 pyramidal neurons

We took confocal z-stack images of GFP-positive, CA3 hippocampal pyramidal neurons, using a Nikon TE2000 C2si laser scanning confocal microscope with a 20× PlanApo objective (0.75 NA), exciting the GFP with a 488 nm laser. Neurons were chosen blind to their genotype or precise topographical position. With the *Thy1*-GFP-M line, we were limited by the overlapping of dendritic branches from neighboring neurons, so we chose to analyze the apical dendrite alone, though the basal dendrites may be accessible for imaging and analysis when using this method with a different transgenic line. During imaging or reconstruction of fluorescent confocal images, if a neuron’s apical dendrite and its neighbors’ dendrites could be distinguished throughout, it was included in the analysis. If there were intersections where it was impossible to make that distinction, we did not include the neuron in the analysis. We only selected neurons that had intact apical arbors. We only selected sections with complete PCP4 staining and no distortions to the section. When taking an image, we positioned the soma at one end of the field-of-view, such that the image captured the entire apical dendrite. The entire thickness of the section was imaged with a step size of 0.7 μm. We imaged a total of 54 neurons from 26 sections from 4 hippocampi for this study: Eleven neurons (from 6 sections) came from a right hippocampus and 17 neurons (from 9 sections) came its left counterpart; 8 neurons (from 4 sections) came from another right hippocampus and 18 neurons (from 7 sections) came from its left counterpart.

### 2.5 Analysis

#### 2.5.1 Dorsoventral angular deviation of primary apical dendrites from transverse plane

We noted that the somata we sampled from were generally found in the middle of the thickness of the section, likely because we only selected neurons with intact apical arbors. Also, in our transverse sections, the primary apical dendrites projected radially away from the soma into the *stratum radiatum*, parallel to the transverse cuts of the section. These two observations suggested that the 3D reconstruction of the apical dendrites is complete. To quantify these observations, we assessed the somatic depth within the section (detailed in 2.5.2 as “depth in section”), and we also assessed the overall dorsoventral angular deviation of the primary apical dendrite from the transverse plane. To assess the overall angular deviation, we sampled confocal images of 25 neurons taken from a left and a right hippocampus taken from one animal. In the confocal images, using Fiji software^25^, we measured the distance between the 3D center of the soma to the endpoint of the primary apical dendrite along the transverse plane (*i.e*., along the xy plane of the image), and we measured the distance between the soma and the endpoint of the primary apical dendrite along the dorsoventral plane (*i.e*., through the z-axis of the confocal image stack). Since these measurements were at right angles from each other, we used them to calculate the angle at which the primary apical dendrite deviated from the transverse plane (in other words, an angle of zero is parallel to the transverse cuts of the section). We also tested the relationship between the dorsoventral positions of the neurons and their angular deviation using linear regression (GraphPad Prism 9).

#### 2.5.2 Topographical positioning: Depth in *stratum pyramidale*, depth in section

The *stratum pyramidale* is distinguishable from the *stratum lucidum* and from the *stratum oriens* in the 20× confocal images we captured. First, we used Fiji software to delineate the border between the *stratum pyramidale* and the *stratum lucidum* and delineated the border between the *stratum pyramidale* and the *stratum oriens*. The “center of *stratum pyramidale*” was arbitrarily defined as the midpoint of those two borders in the radial direction. We then used Fiji software to measure the radial distance between the neuron soma center and the center of the *stratum pyramidale*, with somata superficial to the center of the *stratum pyramidale* recorded as having negative values. We called this distance “depth in *stratum pyramidale*”. We also measured the distance along the z-axis from the top of the section (closest to the coverslip) to the neuron soma center as well as from the bottom of the section (closest to the glass slide) to the neuron soma center to calculate the relative depth of the soma within the section (with 0 meaning that it was adjacent to the coverslip, and 1 meaning that it was adjacent to the glass slide). We called this calculation “depth in section”.

#### 2.5.3 Topographical positioning: DG length, DG width, CA2 length, CA3 length

The granule cell layer of the dentate gyrus is distinguishable in the 4× images we captured. We used FIJI software to measure the length of the granule cell layer, which we called “DG length”. We also measured the distance between the tip of the infrapyramidal blade of the granule cell layer and the tip of the suprapyramidal blade of the granule cell layer, which we called “DG width”.

PCP4 immunohistochemistry allows for the precise identification of CA2 pyramidal neurons, and the labeling of CA2 in this way coincides with the tapering off of the *stratum lucidum*^26,27^. We used the procedure validated by San Antonio *et al*.^28^ (Gaussian blur of 20 μm, 50% local maximal intensity) to determine the immunohistochemical border between CA2 and CA1, as well as between CA2 and CA3. In our case, we used Fiji rather than Adobe Photoshop, utilizing the “Gaussian blur” and “PlotProfile” functions. We then measured the distance (along the superficial border of the *stratum pyramidale*) between the two determined borders, which we called “CA2 length”. We also measured the distance along the *stratum pyramidale* from the border between CA2 and CA3 and the proximal end of the *stratum pyramidale* in CA3, which we called “CA3 length”.

#### 2.5.4 Topographical positioning: Transverse distance from DG, transverse distance from CA2, relative transverse distance

Each cell captured at 20× was registered to the 4× image of its corresponding hippocampal section using Fiji software. We measured the distance (along the superficial border of the *stratum pyramidale*) from the proximal end of the *stratum pyramidale* to the neuron soma center, which we called “transverse distance from DG”. We also measured the distance (along the superficial border of the *stratum pyramidale*) from the border between CA2 and CA3 and the neuron soma center, which we called “transverse distance from CA2”. With these measurements, we calculated the relative distance of the neuron along the *stratum pyramidale* of CA3 (with 0 meaning it is most distal from dentate gyrus, and with 1 meaning it is most proximal to dentate gyrus), which we called “relative transverse distance”. Values between 0 to ⅓ correspond to CA3a (according to the nomenclature coined by Lorente De Nó^29^), values between ⅓ to ⅔ correspond to CA3b, and values between ⅔ to 1 correspond to CA3c.

#### 2.5.5 Topographical positioning: Dorsoventral position

Because we took serial sections, we already knew the dorsoventral (also called “septotemporal”, “long”, or “longitudinal”) position of the neuron relative to the hippocampus analyzed. To record the dorsoventral position of the neuron relative to all hippocampal sections that could be developed with this method, we plotted the DG length, DG width, and CA3 length of each section, which can be predicted by their absolute dorsoventral position in both rats^23^ and mice^30^, regardless of sex or laterality. We assigned an arbitrary number (in this case, zero) to the middle section, and gave positive numbers to more dorsal sections that corresponded to their actual relative distance from the middle section. Sections from other hippocampi were aligned to these sections by aligning their DG length, DG width, and CA3 length to each other. We called this measurement “dorsoventral position”.

#### 2.5.6 Random placement of GFP-positive neurons in CA3

The density of GFP-positive neurons across the tangential extent of CA3 appeared to be random, unlike CA1, where the density of GFP-positive cells clearly increases distally from the dentate gyrus. Within CA3, the likelihood that a particular neuron would be GFP-positive or negative also appeared to be random along the radial and tangential axes, but not along the dorsoventral axis (ventral sections had denser somatic GFP labeling than dorsal sections). To quantify this, we sampled the depth in *stratum pyramidale*, the relative transverse distance, and the dorsoventral position of every 20^th^ GFP-positive CA3 neuron throughout one hippocampus. We counted each GFP-positive neuron in CA3 sequentially, going from CA2 to DG, starting with the most dorsal section, then going from CA2 to DG in the next section, and so on. We captured dorsoventral information from 64 neurons along the dorsoventral position. Of those 64 neurons, we were also able to accurately identify tangential and radial information from 59 and 58 of them, respectively. We used a Kolmogorov-Smirnov test to compare the sampled distribution of values along the sampled range to a theoretical distribution along the same range in which neurons were evenly spaced.

#### 2.5.7 Reconstruction and morphological measurements

We used neuTube software^31^ to generate 3D semi-automated reconstructions from the 20× images we captured. These were saved as standard .swc reconstruction files, which we also inspected using the Simple Neurite Tracer plugin^32^ in Fiji, ensuring that the soma and apical dendrites were categorized correctly by neuTube. Reconstructions are available on the NeuroMorpho.org repository^33,34^. We analyzed the .swc files using L-Measure software^35^. These morphological measurements and any specific settings needed in L-Measure to acquire them are summarized in Supplemental Table 1.

#### 2.5.8 Correlations

We calculated two-tailed, 95% confidence-interval Pearson’s correlation coefficients on all combinations of the 10 position variables to test if any variables had a linear relationship to one another. We chose Pearson’s correlation because there was no *a priori* reason to assume that the data were not going to satisfy the assumptions of the Pearson’s test (*e.g*., samples taken from a normal distribution, random samples, continuous variables, etc.). We eliminated duplicate values to prevent pseudoreplication. To control the false discovery rate (FDR) on these 45 correlations, we applied the method of Benjamini et al.^36^ and set a 10% FDR threshold. We calculated 276 correlation coefficients on all combinations of the 24 morphological variables in the same way. Finally, to compare position variables against morphological variables, we again used the identical procedure to calculate 108 correlation coefficients. We used GraphPad Prism 9 software to calculate statistics and produce graphical representations of statistical calculations.

## 3 Results

### 3.1 GFP and PCP4 expression patterns

In the cerebral cortex and hippocampus, the *Thy1*-GFP-M line almost exclusively targets pyramidal neurons.^20^ We observed that the overall pattern of nearly all GFP-positive somata in the hippocampus proper was restricted to the *stratum pyramidale* of the *cornu Ammonis* and to the granule cell layer of the dentate gyrus (Figure 2). GFP-positive somata were very densely packed in CA1, especially distal to the dentate gyrus, approaching the subiculum. Sparser expression was found in the granule cell layer of the dentate gyrus (Figure 2). The sparsest expression was found in proximal CA1, in CA2, and throughout CA3 (Figure 2). Bright, non-somatic GFP expression was found in the alveus and fimbria (Figure 2), likely representing efferents of GFP-positive somata in the hippocampus proper. Non-somatic expression was also found in the *stratum lucidum* of CA3, likely representing efferent mossy fibers from dentate gyrus. In the *cornu Ammonis*, GFP-positive somata were pyramidal neurons with brightly labeled apical and basal dendrites, and in sparser regions, these dendrites could be distinguished from one another, particularly among apical dendrites. We also observed strong somatic expression of PCP4 in the granule cell layer of the dentate gyrus and strong non-somatic PCP4 expression in the *stratum lucidum* of CA3 (Figure 2). PCP4 expression was also prevalent among CA2 somata, with strong expression extending into their apical and basal dendrites, hence there was also strong PCP4 expression in *stratum radiatum* and *stratum oriens* of CA2 (Figure 2).

**Figure 2.**
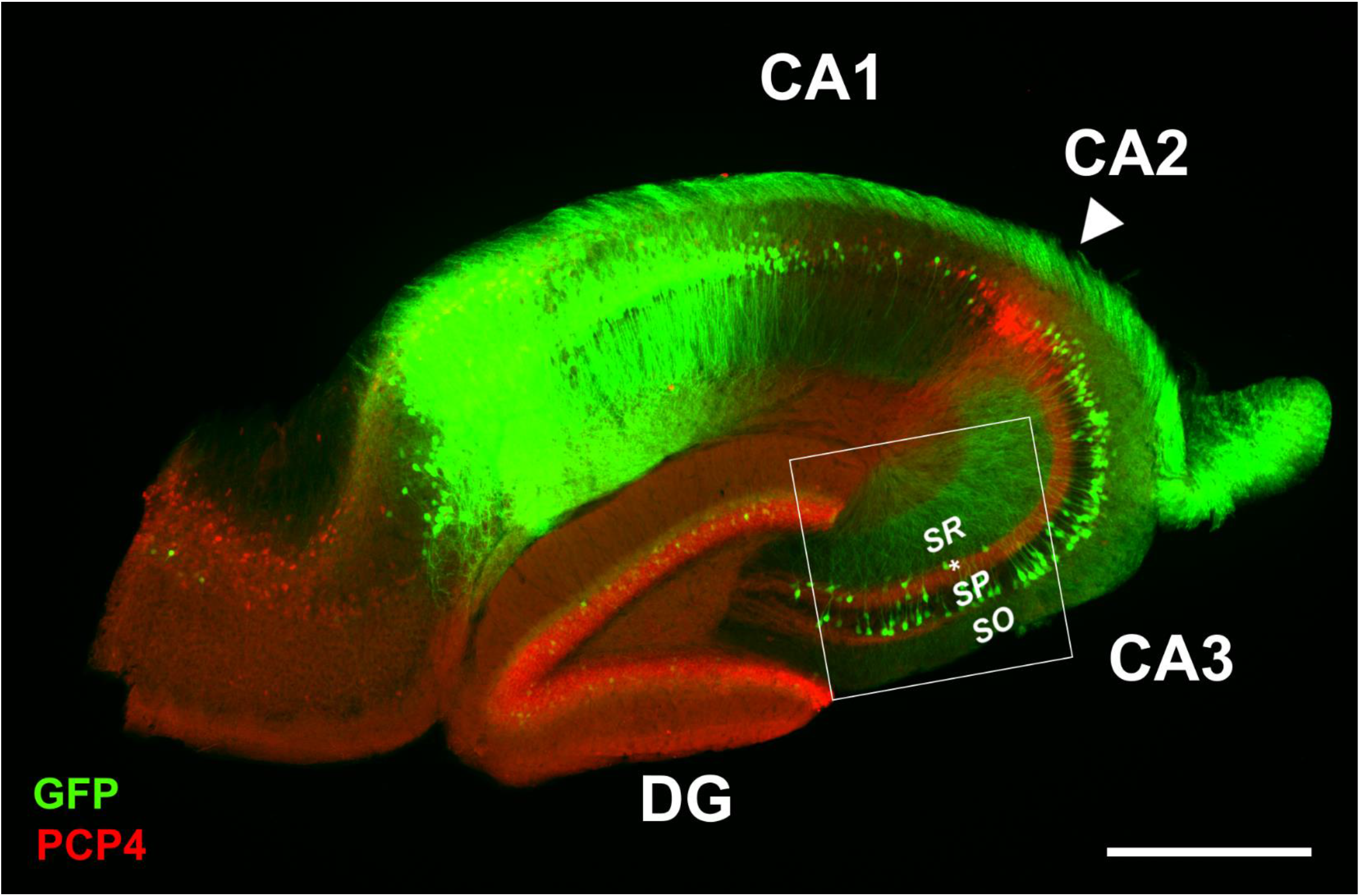
Transverse hippocampal section with fluorescent labeling of PCP4 (red) and GFP (green). DG – dentate gyrus, SR – stratum radiatum, SP – stratum pyramidale, SO – stratum oriens. Arrowhead indicates location of CA2 (PCP4-positive somata), asterisk indicates stratum lucidum (PCP4-positive band of mossy fibers), square indicates area presented in Figure 3. Scale bar: 0.5 mm.

In preparing the hippocampi, we straightened the hippocampus to take exact transverse sections as done previously by other groups.^23,24^ In our case, we hypothesized that the primary dendrite angle does not deviate relative to the transverse plane by using the straightening method. To test this hypothesis, we measured the angle of the primary dendrite of 26 neurons from one hippocampus at various dorsoventral positions. The overall deviation was 6.16° ± 0.64 (Supplemental Table 2). There was no linear correlation between the deviation of the primary dendrite and its dorsoventral position (p > 0.05), as might be expected if there were distortions from straightening. The absolute magnitude of the dorsoventral deviation was also small (4.92 μm ± 0.48; Supplemental Table 2).

The GFP somatic patterning appears mostly unbiased in CA3, in contrast to the clear tangential bias in CA1, where GFP-positive neurons near the subiculum are much more densely concentrated than those close to CA2 (Figure 2). To quantify this observation, we sampled the tangential, dorsoventral, and radial positions of every 20^th^ GFP-positive CA3 neuron from the serial sections of one hippocampus. For each axis (tangential, radial, and dorsoventral), we used a K-S test to compare our empirical data to a theoretic even distribution of neurons along that axis. We found that the neurons are not evenly distributed along the dorsoventral axis (p = 0.0020), with denser patterning in more ventral hippocampus. However, the somatic patterning is not distinguishable from an even distribution along the radial axis (p = 0.6393) or the tangential axis (p = 0.1140).

### 3.2 Topographic data

Each neuron that was 3D reconstructed for apical dendritic morphological analysis was also analyzed for its somatic position along the tangential, radial, and dorsoventral axes in the hippocampus. An example of this is presented in Table 1, which contains topographic information about the three 3D reconstructed neurons presented in Figure 3. In this example, the three neurons have identical values for variables that indicate dorsoventral position (*i.e*., CA2 length, CA3 length, DG length, DG width, dorsoventral position) because they are from the same section. The neurons differ in variables that indicate radial position (*i.e*., depth in *s. pyramidale*) and tangential position (*i.e*., transverse distance from CA2, transverse distance from DG, relative transverse distance; Table 1).

**Table 1.** Topographic and morphological variables for cells presented in Figure 3.

| variable | left<br>(red) | middle<br>(gold) | right<br>(purple) |
| --- | --- | --- | --- |
| CA2 length (μm) | 256 | 256 | 256 |
| CA3 length (μm) | 1176 | 1176 | 1176 |
| DG length (μm) | 890 | 890 | 890 |
| DG width (μm) | 388 | 388 | 388 |
| dorsoventral position (μm) | -300 | -300 | -300 |
| depth in *s. pyramidale* (μm) | -31 | -31 | -30 |
| relative depth in section | 0.51 | 0.34 | 0.63 |
| transverse distance from CA2 (μm) | 1161 | 994 | 824 |
| transverse distance from DG (μm) | 16 | 183 | 352 |
| relative transverse distance | 0.99 | 0.84 | 0.70 |
| total length (μm) | 1508 | 898 | 1036 |
| Euclidean length (μm) | 350 | 295 | 309 |
| path length (μm) | 384 | 364 | 366 |
| primary length (μm) | 57 | 66 | 56 |
| secondary length (μm) | 134 | 287 | 121 |
| tertiary length (μm) | 386 | 225 | 118 |
| quaternary length (μm) | 399 | 146 | 573 |
| overall diameter (μm) | 2.33 | 2.42 | 2.63 |
| total branch points | 7 | 5 | 5 |

**Figure 3.**
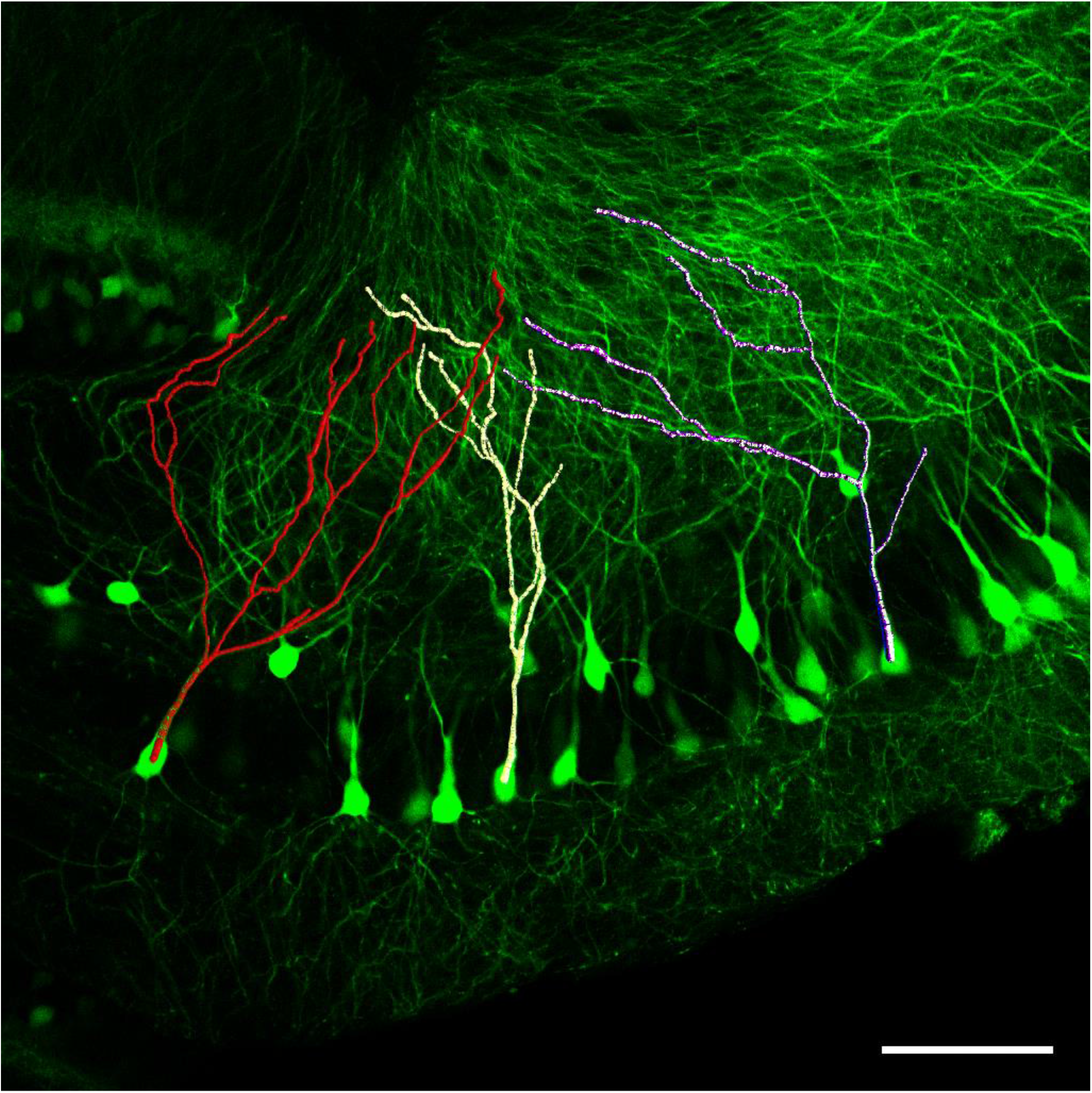
Fluorescent labeling of GFP in CA3 (single optical slice taken from confocal image stack of area delineated in Figure 2). Three 3D-reconstructed neurons included in this study (red, gold, and purple) are overlaid. Topologic and morphological details of the neurons are presented in Table 1. Scale bar: 100 μm.

The overall topographic values for all neurons analyzed in this study are presented in Table 2. Many of the topographic variables appear to be independent: In other words, the dorsoventral, tangential, and radial somatic positions of a neuron do not appear to be related. To investigate this in detail, we tested linear correlations among all pairs of topographic data in each hippocampus. We then tallied the number of hippocampi that had a significant linear correlation between a given pair of topographic variables. Some expected linear relationships emerged. For example, the transverse distance from CA2 and relative transverse distance are linearly correlated in all 4 hippocampi, which is expected because relative transverse distance is calculated by transverse distance from CA2 and from DG (Supplemental Figure 1). Overall, however, most topographic variables have no consistent linear relationship with any other topographic variable (Supplemental Figure 1).

**Table 2.** Topographic variables.

| variable | N | mean | SEM |
| --- | --- | --- | --- |
| CA2 length (μm) | 24 | 224 | 7 |
| CA3 length (μm) | 23 | 1094 | 39 |
| DG length (μm) | 25 | 902 | 18 |
| DG width (μm) | 25 | 346 | 15 |
| dorsoventral position (μm) | 25 | 104 | 73 |
| depth in *s. pyramidale* (μm) | 47 | -28 | 2 |
| relative depth in section | 54 | 0.41 | 0.03 |
| transverse distance from CA2 (μm) | 51 | 612 | 50 |
| transverse distance from DG (μm) | 52 | 499 | 41 |
| relative transverse distance | 50 | 0.54 | 0.04 |

SEM – standard error of the mean

### 3.3 Morphological data

Each 3D reconstructed neuron was analyzed in detail for morphological information about its apical dendrite, including segment length, segment diameter, and branch points (Figure 3; Table 1). The overall morphological values for all apical dendrites analyzed in this study are presented in Table 3. In most hippocampi, almost all morphological variables are independent (*i.e*., pairs of morphological variables are not linearly correlated with one another), except for a linear relationship between Euclidean length and path length (Supplemental Figure 2), which indicates little tortuosity in the dendrites. Because we captured data from two left and two right hippocampi, we were also able to determine that laterality was not an important factor in determining total apical dendritic length (p = 0.0684), though there was a tenuous interaction effect between individual animal and laterality on total apical length (p = 0.0496; Supplemental Table 3).

**Table 3.** Morphological variables.

| variable | N | mean | SEM |
| --- | --- | --- | --- |
| total length ($\mu\text{m}$) | 54 | 1291 | 49 |
| Euclidean length ($\mu\text{m}$) | 54 | 338 | 8 |
| path length ($\mu\text{m}$) | 54 | 388 | 7 |
| primary length ($\mu\text{m}$) | 54 | 61 | 4 |
| secondary length ($\mu\text{m}$) | 54 | 156 | 12 |
| tertiary length ($\mu\text{m}$) | 54 | 292 | 23 |
| quaternary length ($\mu\text{m}$) | 53 | 374 | 27 |
| quinary length ($\mu\text{m}$) | 42 | 314 | 24 |
| senary length ($\mu\text{m}$) | 26 | 281 | 44 |
| septenary length ($\mu\text{m}$) | 11 | 174 | 26 |
| overall diameter ($\mu\text{m}$) | 54 | 2.27 | 0.03 |
| primary diameter ($\mu\text{m}$) | 54 | 3.81 | 0.09 |
| secondary diameter ($\mu\text{m}$) | 54 | 2.51 | 0.05 |
| tertiary diameter ($\mu\text{m}$) | 53 | 2.27 | 0.04 |
| quaternary diameter ($\mu\text{m}$) | 53 | 2.21 | 0.03 |
| quinary diameter ($\mu\text{m}$) | 42 | 2.15 | 0.04 |
| senary diameter ($\mu\text{m}$) | 26 | 2.12 | 0.05 |
| septenary diameter ($\mu\text{m}$) | 11 | 2.21 | 0.07 |
| total branch points | 54 | 7.4 | 0.4 |
| tertiary branches | 54 | 3.6 | 0.1 |
| quaternary branches | 53 | 4.1 | 0.2 |
| quinary branches | 42 | 4.0 | 0.3 |
| senary branches | 26 | 3.4 | 0.4 |
| septenary branches | 11 | 2.7 | 0.3 |

SEM – standard error of the mean

Finally, we wondered whether the 3D somatic position of a neuron was correlated with its 3D apical dendritic morphology in any way, but we found no linear correlation in any topographic-morphological pair of variables in our data.

## 4 Discussion

We developed a method that simultaneously records the detailed 3D position of a neuron (with 10 topological variables) as well as its detailed 3D morphology (with 24 morphological variables). The method is designed to be compatible with a wide variety of reporter lines and immunohistochemical techniques. In developing the method, we found that most topographic and morphological variables of CA3 pyramidal neurons were independent from each other. We further found that CA3 apical dendritic morphology of pyramidal neurons does not change significantly along the dorsoventral, proximodistal, or radial axes in the parts of the hippocampus that we sampled, which agrees with prior work. For example, Malik *et al*.^23^ showed that there is little variability in apical dendritic length along most of the middle of the dorsoventral axis (their Figure 5C). Ishizuka *et al*.^37^, which took the middle third of the dorsoventral length of CA3, reported no significant changes in CA3 apical morphology along the proximodistal axis in the *stratum radiatum* (their Figure 9). An area and dendritic compartment in CA3 that does not change due to somatic position reduces confounds for investigators who are investigating the effect of a particular gene or environmental treatment on CA3 dendritic morphology.

Using the *Thy1*-GFP-M line allows for sparse GFP labeling of pyramidal neurons in CA3 in a seemingly random fashion. We found that the somatic GFP labeling pattern is unbiased in the tangential and radial direction, but also found that the somatic GFP labeling is denser in more ventral sections. However, we did not find a linear relationship between the dorsoventral positions of CA3 neurons and their apical dendritic morphology, so there is no evidence to believe that the non-random distribution of GFP-positive somata found along the dorsoventral axis affects the data collection or analysis of this method in any important way.

We straightened the hippocampus to take exact transverse sections^23,24^. Gaarskjaer^24^ discussed potential distortions with and without straightening in relation to hippocampal cytoarchitecture, but to our knowledge, potential distortions with and without straightening on dendritic morphology have not been investigated until now. In our case, we straightened all hippocampi, so we obviously cannot conclude about the potential for distorted dendritic morphology if the hippocampus is not straightened prior to sectioning. However, in our data, there was no linear correlation between the angle of the primary dendrite and its dorsoventral position (p > 0.05), suggesting that there are no obvious distortions from straightening. Straightening the hippocampus to section the hippocampus exactly perpendicular to its dorsoventral axis ensures that the position of the neuron is accurately recorded along the tangential, radial, and dorsoventral axes.

Several limitations apply to this study:

- *Number of neurons*: With the Thy1-GFP-M line, 17-18 neurons seemed to be an upper limit to the number of CA3 neurons that could be analyzed per hippocampus. With this sample size, linear relationships with an r^2^ of 0.40 or higher can be detected with 80% power in a single hippocampus. However, we believe this limitation is based on the line we chose (*Thy1*-GFP-M) rather than the method overall. The number of analyzed neurons per hippocampus could be increased by choosing a different fluorescent line or choosing to analyze only primary/secondary branch orders, which are much easier to distinguish from neighbors.
- *Dendritic compartments analyzed*: We were only able to reconstruct the apical dendrites of CA3 pyramidal neurons using the *Thy1*-GFP-M line, as basal dendrites from adjacent fluorescent neurons would intersect too much, making them indistinguishable from each other. Apical dendrites are approximately 49-58% of the total dendritic length^37^. Again, we believe this limitation is based on the fluorescent line, so an alternative approach would be to adapt this method to another transgenic labeling strategy.
- *Cell type*: The *Thy1*-GFP-M line almost exclusively targets pyramidal neurons of cerebral cortex and hippocampus^20^ and nearly all targeted cells in CA3 were restricted to *stratum pyramidale* and appeared to have pyramidal morphology. Instead of the *Thy1*-GFP-M line, other transgenic lines could be combined with this method to label other cell types, such as interneurons.
- *Perfusion time*: We transcardially perfused mice with saline (PBS) followed by fixative (4% PFA) after 5-10 minutes. An electron microscopy study has shown alterations to the postsynaptic density and somatic size with a delayed introduction of fixative after the opening of the thoracic wall.^22^ Our study used different perfusates (4% PFA), temperatures (ice cold), duration of fixative perfusion (5-10 minutes), and post-fixation times (24 h). Nonetheless, investigators—particularly those studying the postsynaptic density or somatic size—are urged to consider their perfusion fixation parameters carefully.
- *Section thickness*: In this method, choosing the section thickness is a balance between achieving higher resolution of dorsoventral topographic information (by taking thinner sections) and higher resolution of the overall dendritic morphology (by taking thicker sections), so the optimal thickness will depend on the study objectives. The section thickness in this study (100 μm) was sufficient for accurately reconstructing apical dendrites (otherwise, we should have seen an obvious effect of “depth in section” on the morphology of the neuron if this were a significant problem).
- *Sex as a biological variable*: Sampling from male and female mice allows investigators to assess the effect of sex as a biological variable on their morphological studies. This would be tested similarly to how we studied the potential effect of laterality on dendritic morphology. We could not test for this in our analysis, as our sample happened to include only females. Furthermore, we did not attempt to measure their estrus cycle at this young age (postnatal day 30).
- *Laterality as a biological variable*: We sampled from two left and two right hemispheres. While we could test for laterality differences in the dendritic morphology, these comparisons were statistically weaker compared to the other data presented here, due to clustering.
- *Age*: We only used postnatal day 30 animals in this study that were not visibly abnormal in weight (as the runt of the litter might be). We used the *Thy1*-GFP-M line, which was first presented by Feng *et al*. (2000) along with many other similar lines.^20^ Investigators using the *Thy1*-YFP-H line found that YFP in YFP-positive neurons at 1 month of age is stable for months, but that the density of YFP-positive neurons gradually increases from 1 month of age to 4-6 months of age.^38^ Thus, if the YFP-H line is any indication, we might expect the density of GFP+ neurons to increase gradually from the 1-month mark as well. Nonetheless, YFP+ neurons are still relatively sparse at 4-6 months of age, enabling detailed morphological analysis^21,38^, so we would not expect this increase to significantly hamper investigations in the GFP-M line.
- *Nonlinear relationships*: Our statistical analysis was restricted to testing linear relationships between pairs of variables, so we cannot exclude the possibility that variable pairs are related in some nonlinear way.

The main benefit of the method is that native genetic fluorescent labeling can be leveraged for 3D morphological reconstruction and positioning via serial sectioning. This method is therefore most compatible with morphological studies of genetically determined subsets of cells in the mouse that are labeled with fluorescent reporters. Instead of the *Thy1*-GFP-M line, other transgenic lines could be combined with this method to label other cell types, such as interneurons. If other fluorescent lines provide can label neurons sparsely enough, then other cell types, hippocampal fields, or dendritic compartments could be investigated (including neurons in CA1 or even dentate gyrus). We focused on *Thy1*-GFP-M because it is a very commonly used fluorescent line. The new method presented here produces similar results to diverse approaches for selecting and staining neurons in CA3, such as neurobiotin fills^37^, horseradish peroxidase fills^23^, and Golgi-Cox staining.^30^

Another benefit to this method is that it allows for determining the exact 3D somatic position of reconstructed neurons. In other words, we can take serial sections to determine 3D somatic position *and also* 3D reconstruct neurons in those same sections. In other morphological studies, topographic data have been approximated by using serial sections that are different from the sections in which individual neurons were reconstructed^23,30^: individual reconstructed neurons are later “mapped” to the topographic information gleaned from the serial sections. This strategy improves upon a more approximate method, in which sections are mapped to a mouse brain atlas (*e.g*., Paxinos and Franklin^39^ or the Allen Mouse Brain Atlas^40^). To our knowledge, our method is unique because we assigned reconstructed neurons directly to an exact 3D position without the need for a separate map.

In conclusion, we were able to leverage native genetic fluorescent labeling to create 3D morphological reconstructions while preserving information about the precise 3D somatic position of the neuron, using simple laboratory techniques and a commonly used transgenic fluorescent line. Increased access to detailed topographic and morphological information is an important objective for eliminating confounds and increasing rigor and reproducibility in morphological studies of hippocampal pyramidal neurons.

## Contributors

CH: conceptualization, data curation, formal analysis, investigation, methodology, project administration, supervision, validation, visualization, writing – original draft, writing – review & editing. KB: conceptualization, investigation, methodology, project administration, supervision. CB, CD, AK, BS, HR, HC, EV, MF: investigation, methodology. GV: conceptualization, data curation, formal analysis, funding acquisition, investigation, methodology, project administration, resources, supervision, validation, visualization, writing – original draft, writing – review & editing.

## Acknowledgements

We are exceedingly grateful to Mses. Stephanie Atkins and Sarah Keegan for their superlative animal care.

