## Supplemental Information for "Simultaneous 3D Cellular Positioning and Apical Dendritic Morphology of Transgenic Fluorescent Mouse CA3 Hippocampal Pyramidal Neurons"

**Supplemental Table 1. Morphological measurement settings in L-Measure**

| Type | Name | Description | L-Measure: "Specificity" tab settings | L-Measure: "Function" tab settings | L-Measure: "Output" tab settings |
|---|---|---|---|---|---|
| Length | Total length | The sum of the total length of dendrites of all analyzed apical branch orders | Type = 4 (specifies apical dendrites); Branch_Order < 7 (primary through septenary dendrites) | Length | Total_Sum |
| Length | Euclidean length | The longest distance from the soma to the tip of an apical dendrite, measured "as the crow flies" | Type = 4 | EucDistance | Maximum |
| Length | Path length | The longest distance from the soma to the tip of an apical dendrite, measured along the dendrite | Type = 4 | PathDistance | Maximum |
| Length | Primary length | The length of the primary apical dendrite | Type = 4; Branch_Order = 0 | Length | Total_Sum |
| Length | Secondary length | The total length of the secondary apical dendrites | Type = 4; Branch_Order = 1 | Length | Total_Sum |
| Length | Tertiary length | The total length of the tertiary apical dendrites | Type = 4; Branch_Order = 2 | Length | Total_Sum |
| Length | Quaternary length | The total length of the quaternary apical dendrites | Type = 4; Branch_Order = 3 | Length | Total_Sum |
| Length | Quinary length | The total length of the quinary apical dendrites | Type = 4; Branch_Order = 4 | Length | Total_Sum |
| Length | Senary length | The total length of the senary apical dendrites | Type = 4; Branch_Order = 5 | Length | Total_Sum |
| Length | Septenary length | The total length of the septenary apical dendrites | Type = 4; Branch_Order = 6 | Length | Total_Sum |
| Diameter | Overall diameter | The average diameter of the entire apical dendrite | Type = 4 | Diameter | Average |
| Diameter | Primary diameter | The average diameter of the primary apical dendrite | Type = 4; Branch_Order = 0 | Diameter | Average |

| | | | | | |
|---|---|---|---|---|---|
| Diameter | Secondary diameter | The average diameter of the secondary apical dendrites | Type = 4; Branch_Order = 1 | Diameter | Average |
| Diameter | Tertiary diameter | The average diameter of the tertiary apical dendrites | Type = 4; Branch_Order = 2 | Diameter | Average |
| Diameter | Quaternary diameter | The average diameter of the quaternary apical dendrites | Type = 4; Branch_Order = 3 | Diameter | Average |
| Diameter | Quinary diameter | The average diameter of the quinary apical dendrites | Type = 4; Branch_Order = 4 | Diameter | Average |
| Diameter | Senary diameter | The average diameter of the senary apical dendrites | Type = 4; Branch_Order = 5 | Diameter | Average |
| Diameter | Septenary diameter | The average diameter of the septenary apical dendrites | Type = 4; Branch_Order = 6 | Diameter | Average |
| Branching | Total branch points | The total number of branch points on the entire apical dendrite | Type = 4 | N_bifs | Total_Sum |
| Branching | Tertiary branches | The total number of branches that are tertiary dendrites | Type = 4; Branch_Order = 2 | N_branch | Total_Sum |
| Branching | Quaternary branches | The total number of branches that are quaternary dendrites | Type = 4; Branch_Order = 3 | N_branch | Total_Sum |
| Branching | Quinary branches | The total number of branches that are quinary dendrites | Type = 4; Branch_Order = 4 | N_branch | Total_Sum |
| Branching | Senary branches | The total number of branches that are senary dendrites | Type = 4; Branch_Order = 5 | N_branch | Total_Sum |
| Branching | Septenary branches | The total number of branches that are septenary dendrites | Type = 4; Branch_Order = 6 | N_branch | Total_Sum |

**Supplemental Table 2A. Somatic depth in section and primary apical dendritic angle.**

| **Measurement** | **Mean ± SEM** |
|---|---|
| A: Distance between soma and distal endpoint of primary apical dendrite on the transverse plane | 48.76 ± 2.408 µm |
| B: Distance between soma and distal endpoint of primary apical dendrite on the dorsoventral plane | 4.920 ± 0.4828 µm |
| C: Dorsoventral angular deviation of primary apical dendrite from transverse plane | 6.160 ± 0.6750 ° |

N = 25 cells from one left and one right hippocampus from one animal.

**Supplemental Table 2B. Individual data points of measurements given in Supplemental Table 2A.**
Letters A-C represent the same measurements given in Supplemental Table 2A.

| **A** (µm) | **B** (µm) | **C** (°) |
|---|---|---|
| 63 | 4 | 4 |
| 55 | 4 | 4 |
| 59 | 1 | 1 |
| 51 | 4 | 4 |
| 42 | 5 | 6 |
| 39 | 3 | 5 |
| 56 | 4 | 4 |
| 49 | 6 | 8 |
| 54 | 3 | 3 |
| 43 | 2 | 3 |
| 34 | 3 | 5 |
| 35 | 7 | 11 |
| 47 | 7 | 8 |
| 69 | 6 | 5 |
| 47 | 4 | 4 |
| 53 | 11 | 11 |
| 24 | 4 | 10 |
| 41 | 5 | 7 |
| 59 | 5 | 5 |
| 61 | 3 | 3 |
| 76 | 4 | 3 |
| 42 | 9 | 12 |
| 33 | 6 | 11 |
| 41 | 3 | 4 |
| 46 | 10 | 13 |

**Supplemental Table 3. Two-way ANOVA of total length (laterality, animal)**

| **factor 1** | **factor 2** | **factor tested** | **DFn** | **DFd** | **F** | **p-value** |
|---|---|---|---|---|---|---|
| laterality | animal | laterality | 1 | 50 | 3.470 | 0.0684 |
| laterality | animal | animal | 1 | 50 | 0.009 | 0.9237 |
| laterality | animal | interaction | 1 | 50 | 4.049 | 0.0496 |

DFn – degrees of freedom numerator; DFd – degrees of freedom denominator.

**Supplemental Figure 1.**

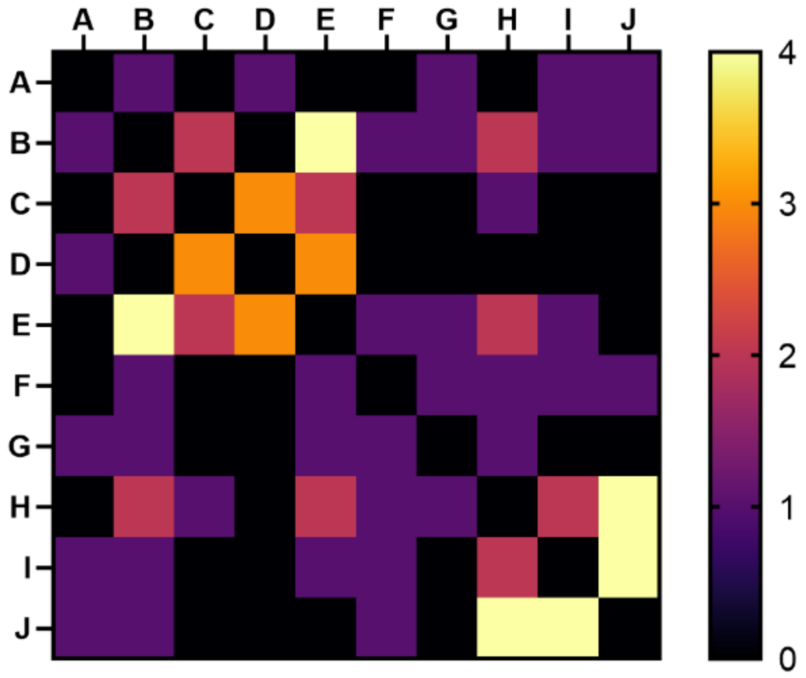


**"Consensus" matrix showing the number of hippocampi with p-values under the 10% FDR threshold for pairs of topographic variables.** Key: **A**, CA2 length (µm); **B**, CA3 length (µm); **C**, DG length (µm); **D**, DG width (µm); **E**, dorsoventral position (µm); **F**, depth in *s. pyramidale*; **G**, depth in section; **H**, transverse distance from CA2 (µm); **I**, transverse distance from DG (µm); **J**, relative transverse distance.

**Supplemental Figure 2.**

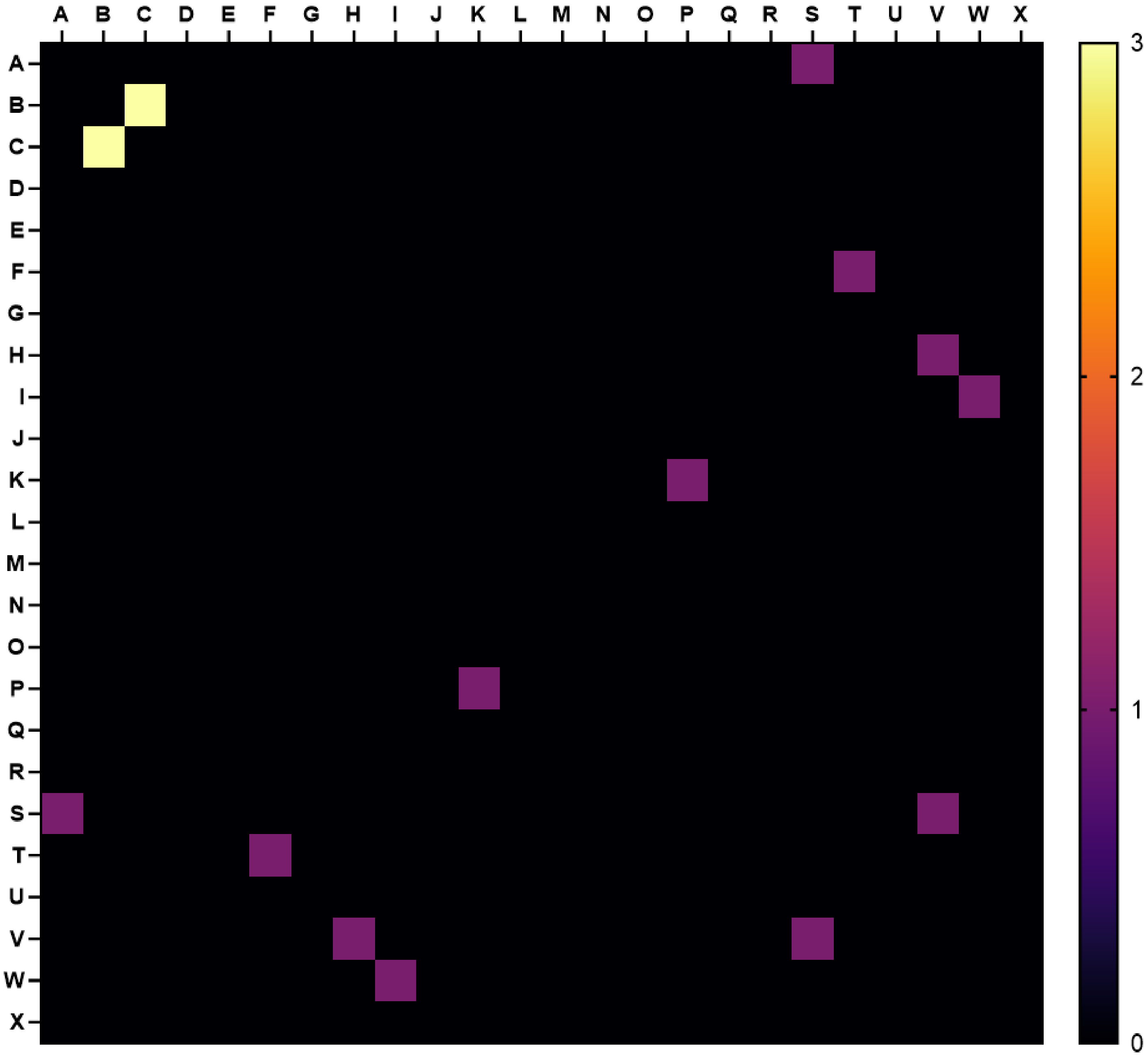


**“Consensus” matrix showing the number of hippocampi with p-values under the 10% FDR threshold for pairs of morphological variables** (hippocampus 1 $p < 0.00044$, hippocampus 2 $p < 0.00101$, hippocampus 3 $p < 0.00036$, hippocampus 4 $p < 0.00177$). Key: **A**, total length (µm); **B**, Euclidean length (µm); **C**, path length (µm); **D**, primary length (µm); **E**, secondary length (µm); **F**, tertiary length (µm); **G**, quaternary length (µm); **H**, quinary length (µm); **I**, senary length (µm); **J**, septenary length (µm); **K**, overall diameter (µm); **L**, primary diameter (µm); **M**, secondary diameter (µm); **N**, tertiary diameter (µm); **O**, quaternary diameter (µm); **P**, quinary diameter (µm); **Q**, senary diameter (µm); **R**, septenary diameter (µm); **S**, total branch points; **T**, tertiary branches; **U**, quaternary branches; **V**, quinary branches; **W**, senary branches; **X**, septenary branches
